# H2A.Z is dispensable for both basal and activated transcription in post-mitotic mouse muscles

**DOI:** 10.1101/823526

**Authors:** Edwige Belotti, Nicolas Lacoste, Thomas Simonet, Christophe Papin, Kiran Padmanabhan, Lorrie Ramos, Defne Dalkara, Isabella Scionti, Ali Hamiche, Stefan Dimitrov, Laurent Schaeffer

## Abstract

The histone variant H2A.Z is enriched in nucleosomes surrounding the transcription start site of active promoters, suggesting that it might be implicated in transcription. It is also required during mitosis. However, evidences obtained so far mainly rely on correlative evidences obtained in actively dividing cells. We have defined a paradigm in which cell cycle cannot interfere with H2A.Z transcriptional studies by developing an *in vivo* systems to invalidate H2A.Z in terminally differentiated post-mitotic muscle cells to dissociate its role during transcription from its role during mitosis. ChIP-seq, RNA-seq and ATAC-seq experiments performed on H2A.Z KO post-mitotic muscle cells show that this histone variant is neither required to maintain nor to activate transcription. Altogether, this study provides *in vivo* evidence that in the absence of mitosis H2A.Z is dispensable for transcription and that the enrichment of H2A.Z on active promoters is rather a marker than an actor of transcriptional activity.

## INTRODUCTION

Histone variants are non-allelic isoforms of conventional histones^1,2^. Each histone has histone variants even H4 but only in hominids^1,3,4^. Histone variants differ in their primary sequence, transcriptional regulation and timing of expression during the cell cycle compared to conventional histones^1–3^. Incorporation of histone variants can confer novel structural properties to nucleosomes and change the functional chromatin landscape^3,5–7^. The current view is that histone variants are involved in all aspects of DNA metabolism including transcription, replication and repair^1^. They are likely implicated in numerous diseases, particularly in cancer (for review see^8,9,10^).

The H2A family of histone variants is one of the richest. In addition to conventional H2A, it comprises at least the histone variants macroH2A, H2A.X, H2A.Z and H2A.Bbd^11^.

In mammals, the H2A.Z variant comprises two proteins: H2A.Z-1 and H2A.Z-2^12,13^. They are encoded by two distinct genes located on two different chromosomes (Chr3 and Chr11, respectively) and they differ by only three amino-acids^12^. H2A.Z is an essential protein in mammals^14^. ANP32E and YL1 are part of large protein complexes and were identified as H2A.Z specific chaperones, responsible for either its eviction or its deposition in chromatin^15,16^. In vertebrates, genome-wide studies have shown that H2A.Z is enriched at active promoters^17–19^, in facultative heterochromatin^20^ and at centromeres^21,22^. Interestingly, +1 and −1 nucleosomes surrounding the Transcription Start Site (TSS) are enriched in H2A.Z in proportions correlated with the strength of transcription^18,23^. H2A.Z is also preferentially bound to other regulatory elements such as enhancers and CTCF-binding sites, which mark insulator sites in the genome^15,17,24^ and more generally in chromatin regions where histones have a high turnover. From all these observations, a consensus emerged stipulating that H2A.Z was a key player in the control of transcription^17,18,25^.

However, the available data on the link between H2A.Z-containing nucleosomes and transcription remain mainly correlative and based on studies carried out in cultured cells. The interpretation of such results have to take into account perturbations of replication and cell division that can indirectly affect gene expression. For example, depletion of H2A.Z in vertebrates results in strong mitotic and cytokinesis defects and can activate an apoptotic program subsequently leading to cell death^26^. Thus, in this case the effect of H2A.Z on transcription might be indirect and would be linked with mitotic functions of H2A.Z. It appears that unravelling the exact function of H2A.Z in the control of transcription requires a system in which H2A.Z can be inactivated in cells that do not replicate DNA anymore. Skeletal muscle and the Cre-/loxP system provide such a paradigm.

Transgenic mice, expressing the Cre recombinase under the control of the human α-skeletal actin promoter (HSA-Cre), allow to specifically express the recombinase in postmitotic skeletal muscle cells^27^. Thus, breeding HSA-Cre mice with H2A.Z-1^flox/flox^:H2A.Z-2^flox/flox^ mice specifically inactivates H2A.Z-1 and H2A.Z-2 in postmitotic skeletal muscle cells (hereafter named H2A.Z dKO). These cells provide a convenient tool to analyse *in vivo* the requirement of H2A.Z for transcription maintenance and regulation.

Gene expression in skeletal muscle cells is tightly controlled by innervation. Two days after denervation muscle cells have adopted a new gene expression program involving the activation and the repression of hundreds of genes^28^. The best characterized genes activated upon denervation encode a variety of proteins: transcription factors (MYOD, Myogenin), neurotransmitter receptors (acetylcholine receptor subunits), kinases (MuSK, PAK1), histone deacetylases (HDAC6, HDAC4) and ubiquitin ligases (MURF1, MAFBX, MUSA1)^28–30^. Depriving muscles of their innervation thus provides a convenient mean to evaluate the involvement of H2A.Z in *de novo* regulation of gene expression.

Our RNA-seq and ATAC-seq analysis of innervated and denervated H2A.Z dKO muscles revealed that the absence of H2A.Z affected neither steady state nor activated gene expression.

## RESULTS

### Invalidation of H2A.Z-1 and H2A.Z-2 in MEFs cells

H2A.Z-1^flox/flox^:H2A.Z-2^flox/flox^ mouse line was obtained with the strategy described in figure 1A and B. Briefly, loxP sequences were inserted before exon 2 and after exon 4 of *h2afz* (H2A.Z-1) and before exon 4 and after exon 5 of *h2afv* (H2A.Z-2) using standard recombination strategy. To evaluate the efficacy of our inactivation strategy and the stability of H2A.Z we have first isolated Mouse Embryonic Fibroblasts (MEF) cell lines from the double floxed mouse line. Upon Cre induction using an adenoviral system, *h2afz* and *h2afv* knockdown was was evaluated by RT-qPCR (Fig. 1C), and H2A.Z protein depletion was evaluated by western blotting (Fig. 1D) and immunofluorescence (Fig. 1E). Two days after infection by the adeno-Cre H2A.Z protein was already undetectable in cycling cells. H2A.Z dKO cells did not survive more than 9 days after H2A.Z withdrawal, recapitulating the essential function of H2A.Z^26^. We then used RT-qPCR to evaluate the expression of a set of genes known to be affected by the absence of H2A.Z in mouse cells^25^. As expected, they were misregulated in the absence of H2A.Z (Fig. 1F). We then initiated the study of the role of H2A.Z in postmitotic muscle cells *in vivo* to determine if in absence of replication and cell division the same effects would be observed.

**Figure 1.**
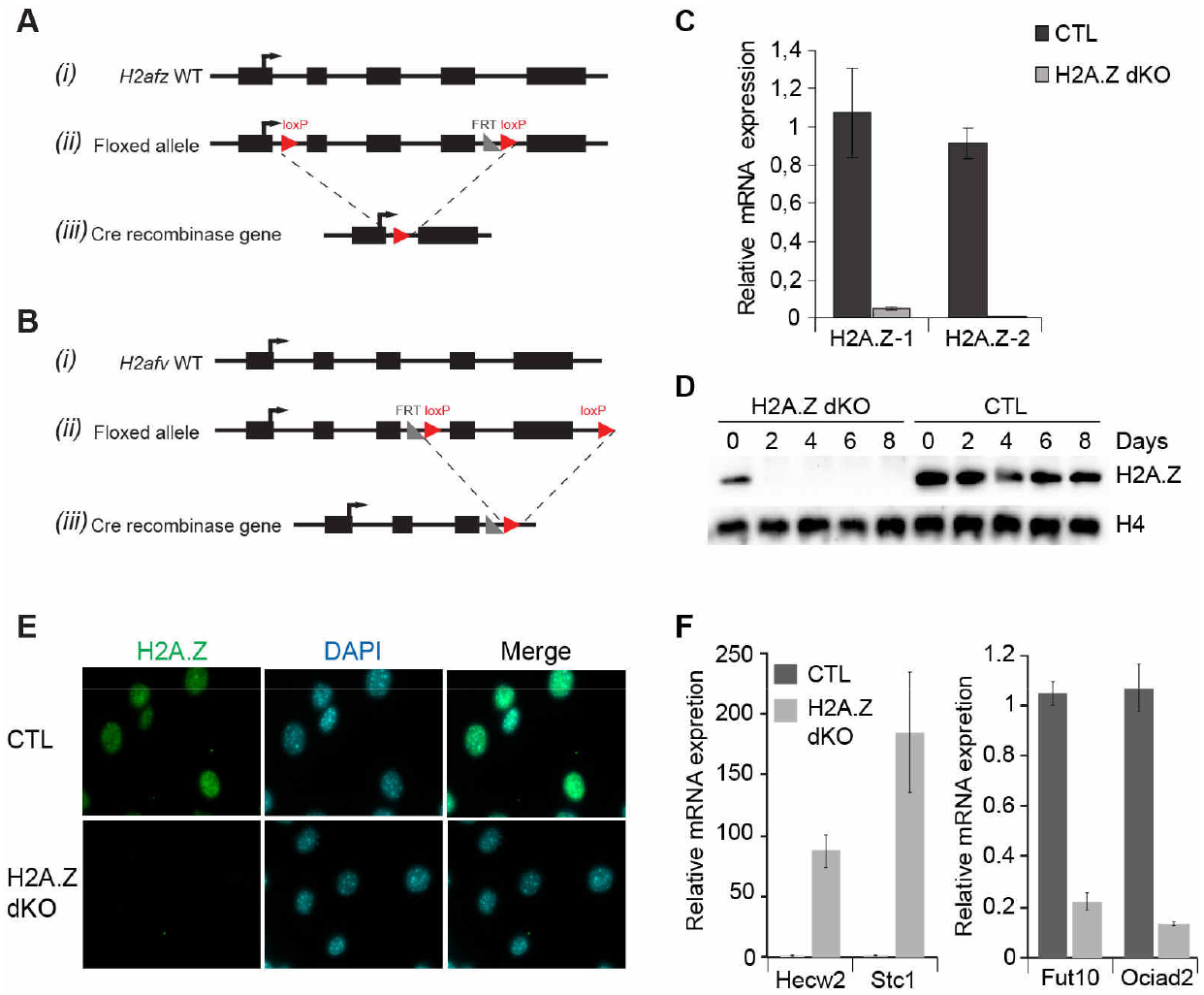
Generation and characterisation of the H2A.Z-1 and H2A.Z-2 models. **(A)** (i) Wild-type *H2afz* gene structure. Black boxes indicate the exons. (ii) The conditional allele of the *H2afz* flox/flox gene. LoxP sites were inserted at the indicated position. (iii) Organization of the H2A.Z-1 knock-out (KO). Exons 2, 3 and 4 are deleted upon expression of Cre recombinase to generate the KO allele *H2afz* (-/-). **(B)** Same as (A), but for the *H2afv* gene. LoxP site was inserted on both ends of exon 4-5. After Cre-recombinase expression, exon 4 and 5 are deleted to create the KO allele *H2afv* (-/-). **(C)** Bar graphs representing the expression level of *H2afz* and *H2afv* genes measured by RT-qPCR analysis in MEFs infected with either control adenovirus or with adenovirus expressing Cre recombinase. **(D)** H2A.Z is efficiently depleted upon infection of MEFs with adenovirus expressing Cre recombinase. Western blot of MEF cells using anti-H2A.Z antibody. H4 was used as loading control. **(E)** Immunofluorescence detection of H2A.Z on proliferating MEFs 2 days after infection. **(E)** RT-qPCR of RNA isolated from CTL and H2A.Z double flox MEFs cells in absence or presence of adeno-Cre virus particles. On the left panel, some up-regulated genes in absence of H2A.Z and on the right panel some down-regulated genes in absence of H2A.Z.

### H2A.Z is enriched at TSS of active genes in mouse skeletal muscle

Currently, no data are available on the distribution of H2A.Z *in vivo* in skeletal muscle and more generally in post-mitotic cells. To analyse H2A.Z genomic distribution in adult skeletal muscle, ChlP-seq was performed on *tibialis anterior* (TA) muscles from 7 weeks old mouse. Results indicated that H2A.Z was enriched around transcriptional start sites (TSS) as in most other cell types^31^ (Fig. 2A). RNA-seq was then performed to compare gene expression data to the ChIP-seq. The results showed that the abundance of H2A.Z was correlated to gene expression (Fig. 2B). Altogether, these results are in agreement with the numerous studies showing that H2A.Z is enriched around the TSS of active genes^15,17,18^. Skeletal muscle thus provides an adequate system to evaluate the requirement of H2A.Z for gene expression in post-mitotic cells.

**Figure 2.**
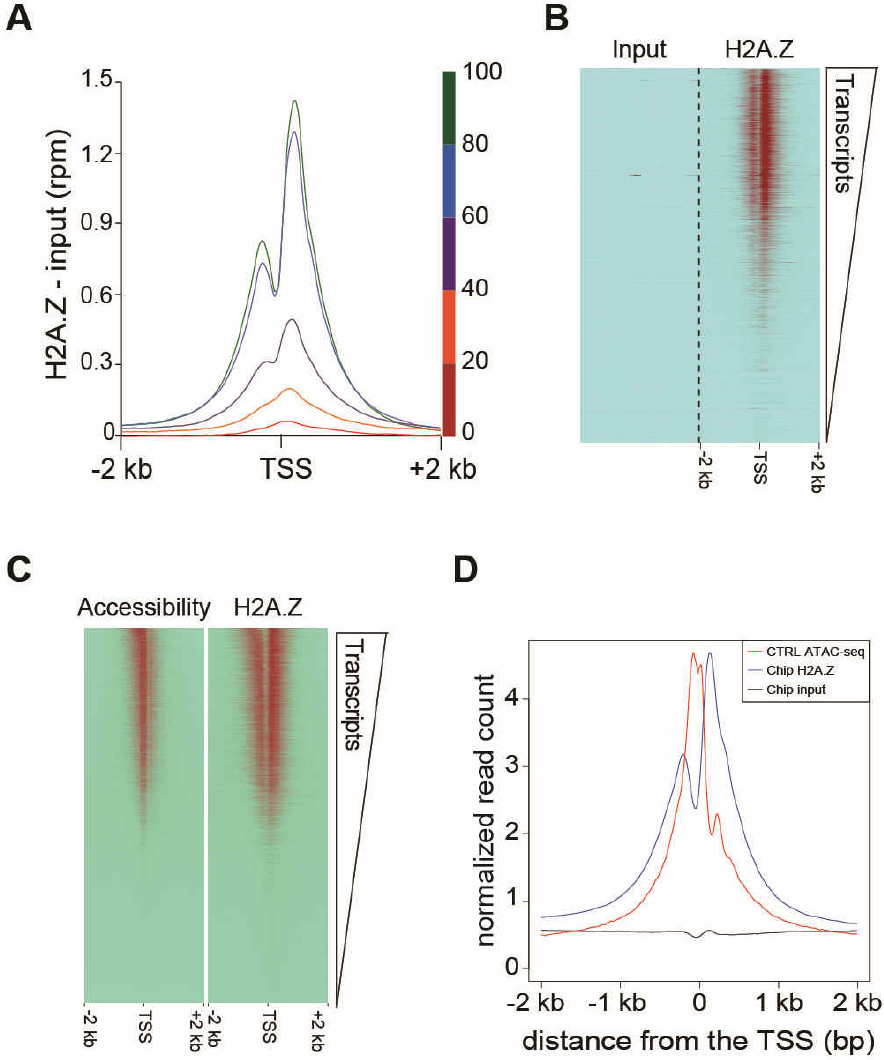
H2A.Z localisation and DNA accessibility at genome-wide level according to the transcription. **(A)** Distribution of H2A.Z relative to the transcription start sites (TSS) (the distinct colors indicate the different levels of transcription) and **(B)** Heat map of input and H2A.Z ChlP-seq enrichments around the transcription start sites (TSSs) ranked according the RNA-seq levels, in muscle skeletal myofibers. **(C)** Heat map of the ATAC-seq signal and H2A.Z Chip-seq enrichment around the transcription start sites (TSSs) ranked according the RNA-seq levels. **(D)** Distribution of H2A.Z and DNA accessibility relative to the TSS.

Using ATAC-seq in control muscles to assess chromatin accessibility at the genome level, we could readily detect a correlation between the presence of H2A.Z and accessible regions of the DNA (Fig. 2C). Since ChIP-seq experiments showed a correlation between the strength of transcription and H2A.Z enrichment at the TSS, we can conclude that in muscle cells, as in any cell type, a correlation exists between transcription strength and chromatin accessibility. ChIP-seq and ATAC-seq alignment further indicated that in muscle fibers, as in other mouse tissues^32^, chromatin was the most accessible between the +1 and the −1 nucleosomes surrounding the TSS (Fig. 2D).

### Invalidation of H2A.Z-1 and H2A.Z-2 in mouse skeletal muscle fibers

The H2A.Z-1^flox/flox^:H2A.Z-2^flox/flox^ mouse line was crossed with mice expressing the Cre recombinase under the control of the HSA promoter that specifically expresses the Cre recombinase in post-mitotic skeletal muscle cells^27^. In HSA-Cre mice, the expression of the Cre recombinase is initiated in the second half of embryonic development and reaches its plateau of expression around birth^33^. The depletion of both H2A.Z in the TA muscles of 7-8 weeks old mice was demonstrated by RT-qPCR (Fig. 3B) and western blotting (Fig. 3C). In both cases, the remaining signal (~15-20 %) reflects the presence of satellite cells and non-muscle cells e.g. fibroblasts, endothelial cells, macrophages^34^. Immunofluorescence experiments were carried out to specifically visualize the decline in H2A.Z content in the nuclei of muscle fibers (Fig. 3A and D). One week after birth H2A.Z levels were already significantly decreased (Fig. 3A) and by post-natal week 4, H2A.Z could not be detected anymore in muscle nuclei. Consistently, 7 weeks after birth no H2A.Z could be detected in muscle nuclei whereas its expression remained unaffected in non-muscle cells (Fig. 3D). For further analysis, muscles were collected seven weeks after birth, thus several weeks after the depletion of H2A.Z (Fig. 3A). Phenotypically, seven weeks old H2A.Z dKO mice were indistinguishable from control mice (CTL). Body weight and muscle histology of these mice were similar to the control ones. Fibrosis, fiber type composition, mitochondria distribution and function were identical in control and H2A.Z dKO muscles (Fig. 4A, see below supplementary Fig. 3). Importantly, the distribution of fiber size (cross-sectional area) was comparable between CTL and H2A.Z dKO muscles (Fig. 4B). In adult muscles, the loss of muscle fibers is usually compensated by the formation of new muscle fibers from resident adult muscle stem cells. Regenerating muscle fibers are easily detected by the central position of their nuclei since in other fibers the nuclei are located at the periphery, directly beneath the plasma membrane. In healthy muscles the proportion of muscle fibers with central nuclei does not exceed 1%. This value was not increased in H2A.Z dKO muscles, indicating an absence of active regenerative process. Altogether, these data revealed that H2A.Z depletion in skeletal muscle fibers did not induce any detectable histological alteration.

**Figure 3.**
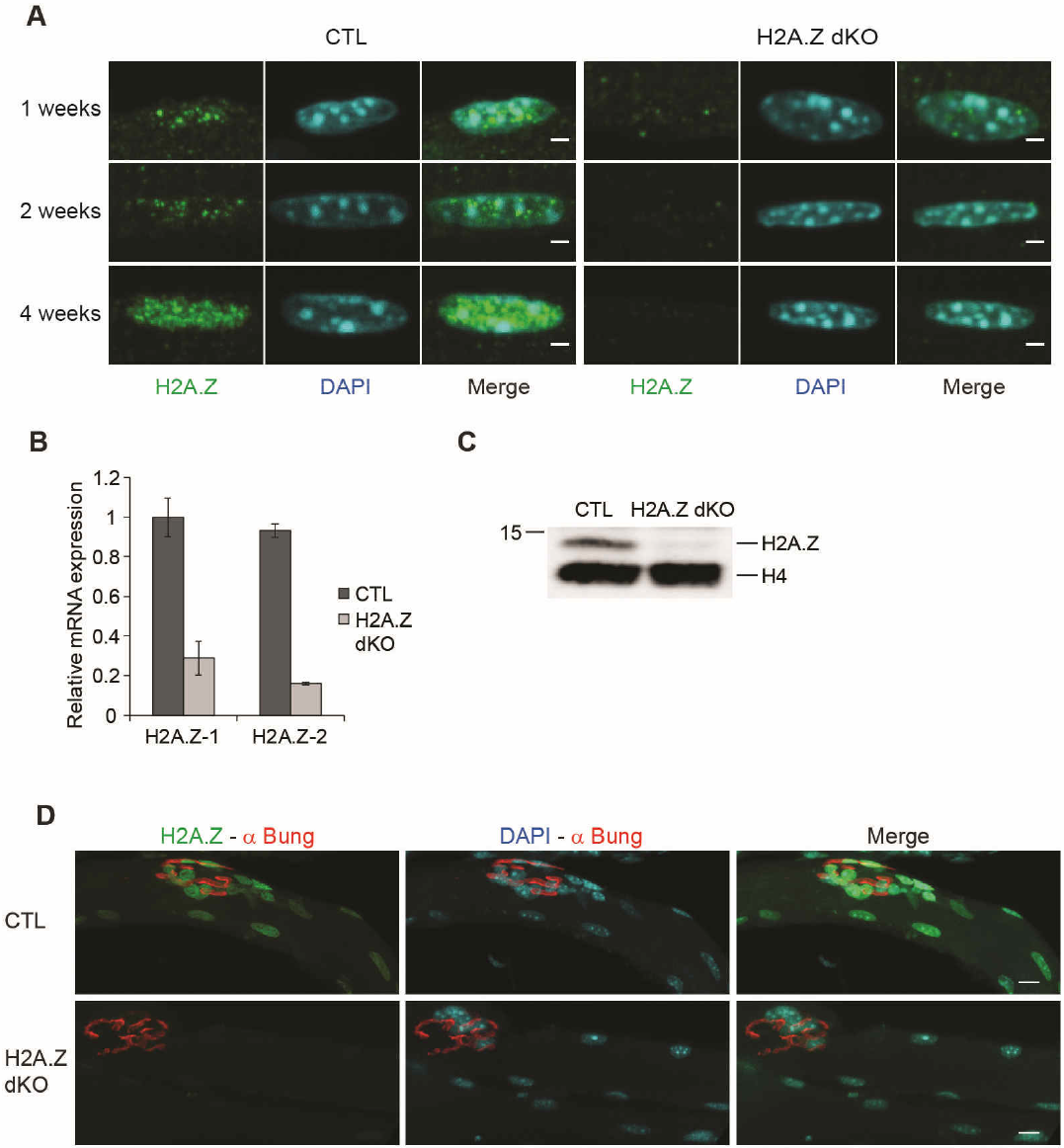
Validation of the H2A.Z cKO mouse model. (A) Immunofluorescence detection of H2A.Z on *tibialis anterior* muscle fibers from 1 week, 2 weeks and 4 weeks old CTL and H2A.Z dKO mice. DAPI was used to stain the DNA (Scale bar 2 μm). **(B)** H2A.Z-1 and H2A.Z-2 RT-qPCR of RNA isolated from the double KO mouse strain on 7 weeks aged mice (n = 3). **(C)** Western blotting of H2A.Z from nuclear extracts of skeletal muscles from CTL and double cKO H2A.Z-1^(-/-)^ x H2A.Z-2^(-/-)^ (H2A.Z dKO) on 7 weeks old mice. H4 is used as loading control. **(C)** Immunofluorescence detection of H2A.Z on EDL muscle fibers from 7 weeks old CTL and H2A.Z dKO mice. α-bungarotoxine is used to stain the neuromuscular junction and DAPI to stain the DNA (Scale bar 10 μm).

**Figure 4.**
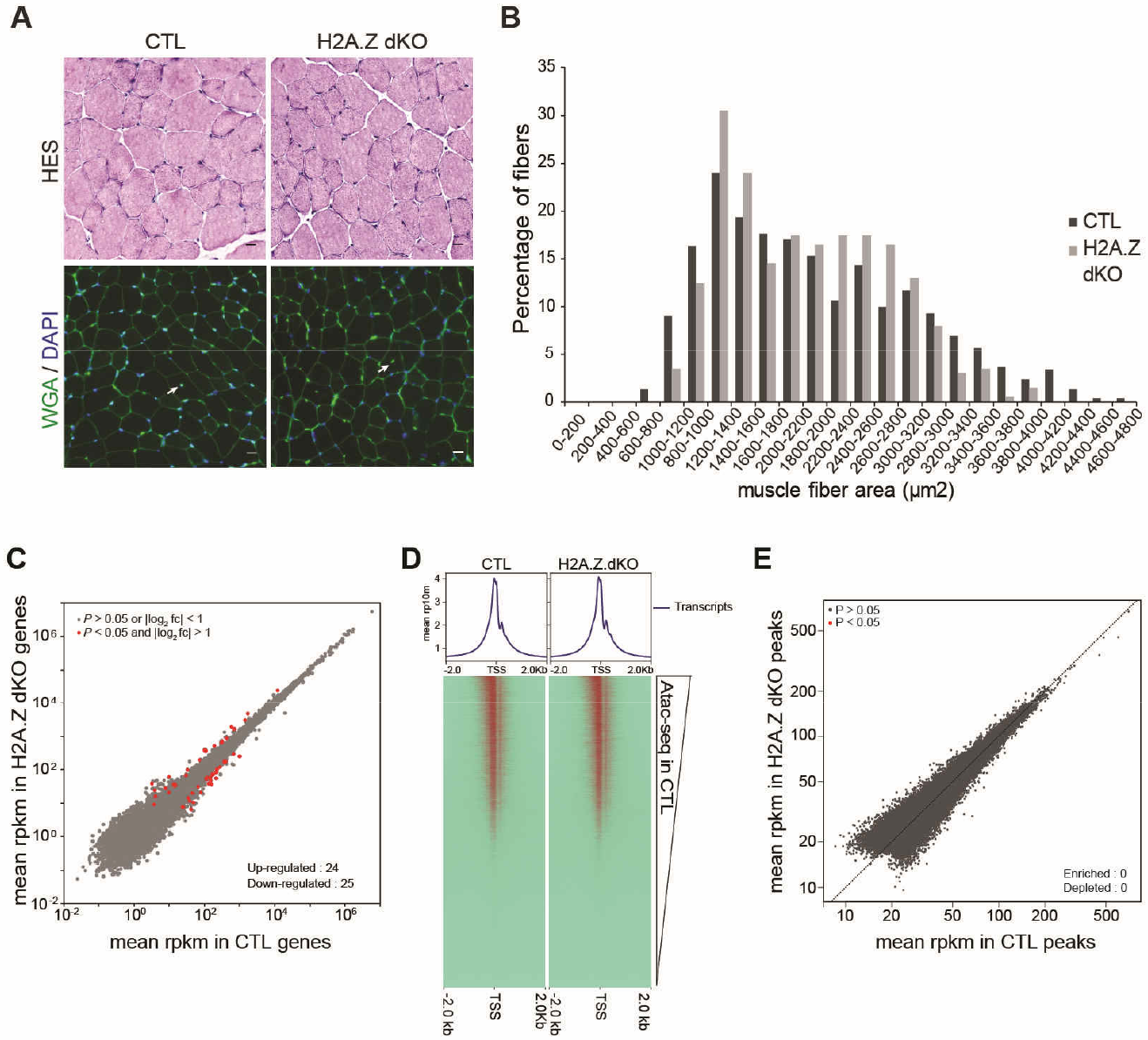
Characterization of the H2A.Z cKO mouse model. **(A)** Histological analysis of *tibialis anterior* muscles by Hematoxyline and Eosin (HE) and Wheat Germ Agglutinin-Lectin (WGA) staining from 7 weeks old CTL and H2A.Z dKO mice (Scale bar 20 μm, arrows indicate centronucleated fibers). **(B)** Fiber size distribution analysed from 200 myofibers of each samples. **(C)** Scatter plots comparing global gene expression levels between CTL and H2A.Z dKO cells in muscle from 7 weeks-old mice. **(D)** Heat map of the ATAC-seq signal enrichment around the transcription start sites (TSSs) in CTL and H2A.Z dKO, the TSSs being ranked according to the CTL level. **(E)** Scatter plot comparing ATAC-seq peaks enrichment in CTL and H2A.Z dKO muscles.

### The absence of H2A.Z perturbs neither steady state gene expression nor the chromatin landscape

To evaluate the impact of H2A.Z inactivation on steady state gene expression, RNA-seq experiment was performed on 7 weeks old CTL and H2A.Z dKO TA muscles. Analysis of results confirmed the inactivation of both H2A.Z isoforms in H2A.Z dKO mice (Supplementary Fig. 1). The transcriptomes of CTL and H2A.Z dKO muscles were almost identical (Fig. 4C). In the absence of H2A.Z, the expression of only 24 and 25 genes was slightly activated or repressed, respectively, and no clustering of these genes was observed (Fig. 4C, Table 1). Therefore, removing H2A.Z from skeletal muscle fibers does not significantly perturb steady state gene expression.

**Table 1.**
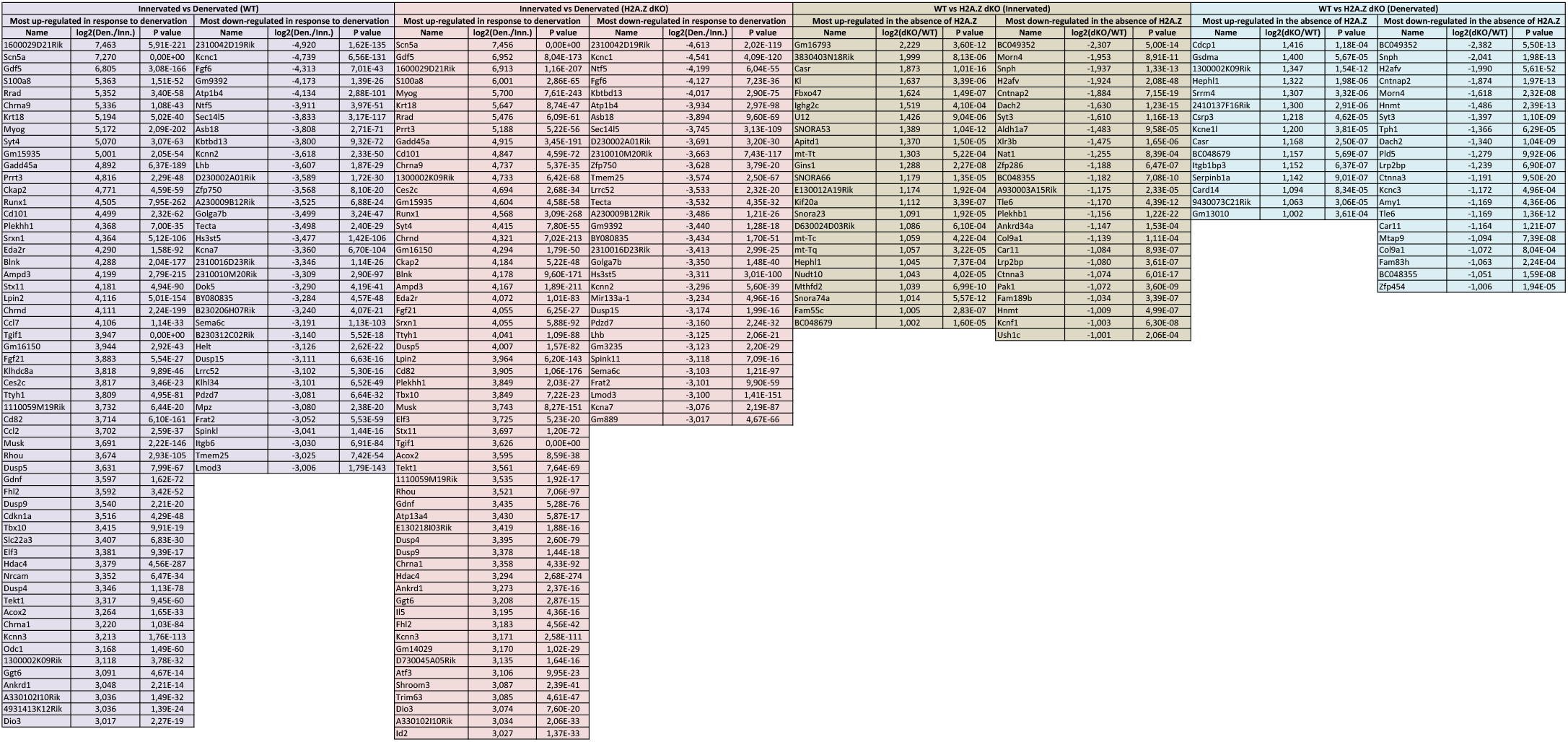
Main down- and up-regulated genes in innervated and denervated CTL and dKO H2A.Z skeletal muscle from 7 week-old mice.

ATAC-seq results sorted by transcriptional strength generated identical patterns in control and H2A.Z dKO muscles (Fig. 4D). This was further confirmed by directly comparing the peaks from each experiment (Fig. 4E). Altogether, these results indicate that in muscle fibers nuclei the absence of H2A.Z did not significantly change gene expression or DNA accessibility.

### The absence of H2A.Z perturbs neither acute gene activation nor repression in mouse skeletal muscle

To evaluate the possibility that H2A.Z would be required to prime transcriptional changes but would be dispensable once promoters were already activated or repressed, we analysed the effect of H2A.Z inactivation upon acute changes in gene expression. For this purpose, seven weeks old CTL and H2A.Z dKO TA muscles were denervated. Denervation is carried out by section of the sciatic nerve of one hind limb. 48h after denervation, TA muscles were collected, the contralateral TA muscles was used as innervated controls. RT-qPCR experiments showed that the strong post-denervation upregulation of *MyoD* and *Myogenin* expression took place normally in the absence of H2A.Z (Supplementary Fig. 2). Note that no histological changes in denervated H2A.Z dKO mice compared to CTL mice have been detected (Supplementary Fig. 3).

To have a global view of the transcriptional response to denervation, RNA-seq was carried out on RNA purified from either innervated or denervated TA muscles. As expected, the expression of *Myogenin, MuSK, Acetylcholine receptor α* and *δ* and *Hdac4* was strongly upregulated (Table 1), thus confirming that muscles responded correctly to denervation. In agreement with available data^35^, the genome-wide transcriptome analysis revealed that upon denervation hundreds of genes were either upregulated (894 genes) or downregulated (944 genes) (Fig. 5A, B). Denervation induced the downregulation of many genes involved in contractility and the upregulation of genes involved in muscle development and inflammation (Fig. 5D). In H2A.Z dKO muscles, 889 genes were upregulated and 876 downregulated. Up- and down-regulated genes were the same in CTL and dKO muscles, indicating that the vast majority of genes is normally regulated by denervation in the absence of H2A.Z (Fig. 5C and Table 1). All together, the depletion of H2A.Z did not significantly affect the genome–wide transcriptional response to denervation (Fig. 5C). We conclude that in post-mitotic skeletal muscle cells, H2A.Z is not required to activate or to repress transcription to adapt to a particular physiological context.

**Figure 5.**
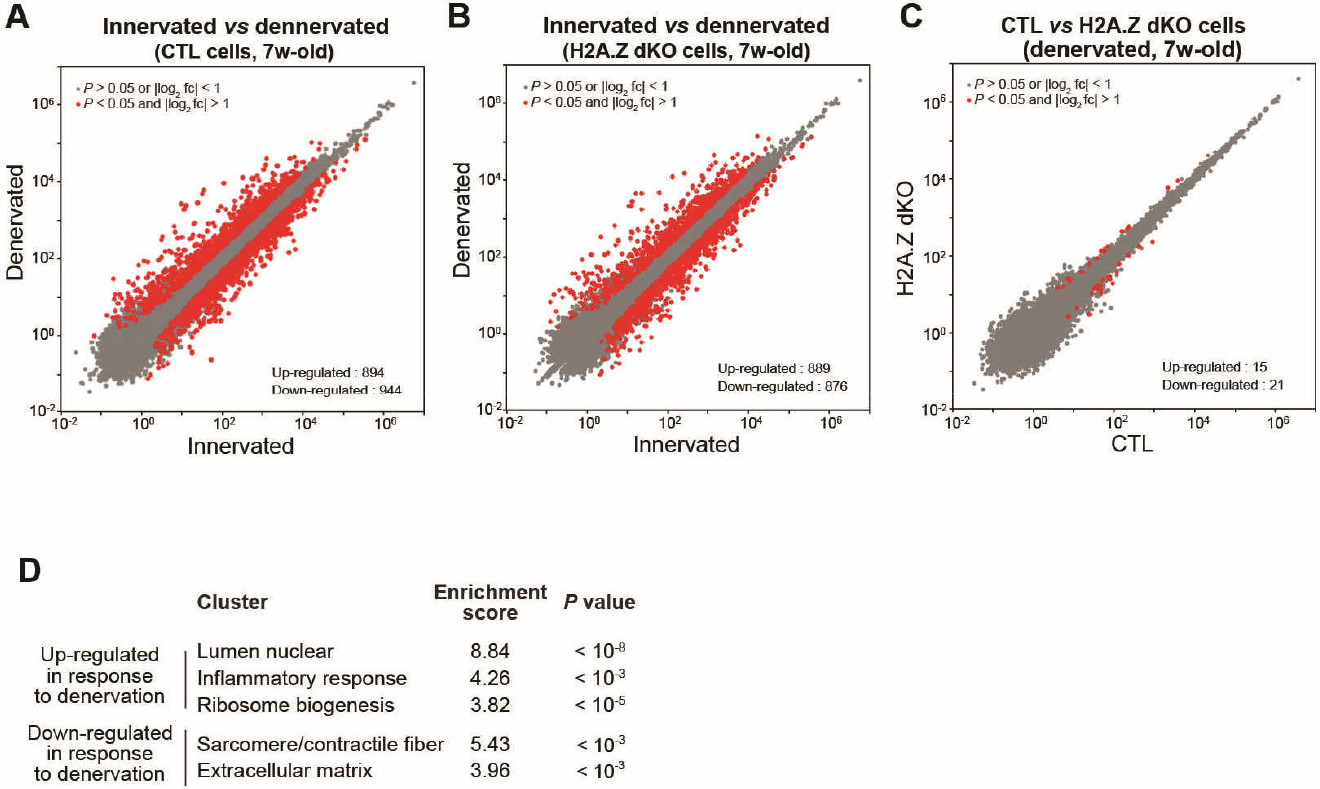
Characterization of the H2A.Z cKO under acute transcriptional response. **(A, B)** Scatter plots comparing global gene expression levels between innervated and denervated cells (in 7 weeks-old mice) in the presence **(A)** or in the absence **(B)** of H2A.Z. **(C)** Scatter plots comparing global gene expression levels 48h after denervation between CTL and H2A.Z dKO cells in muscle cells of 7 weeks-old mice. **(D)** Functional annotation clustering of differentially expressed genes in response to denervation.

### H2A.Z is enriched at DNA repetitive elements

Since H2A.Z is also known to be enriched at pericentromeric regions in the mouse genome, we specifically analysed repetitive sequences that represent a very high proportion of these regions^20,21^. As expected, our ChIP-seq data indicated that numerous families of DNA repetitive elements were enriched in H2A.Z (Fig. 6A). The families that exhibited the higher enrichment are listed in Table 2. Of note, the majority of repeats significantly enriched in H2A.Z (log_2_ enrichment > 0.5 and *P* value < 10^-2^) belonged to the simple repeats and to the endogenous retrovirus classes of repetitive elements. This suggested that H2A.Z might be involved in the control of the expression of these repetitive DNA elements. To test this hypothesis, we analysed how the absence of H2A.Z affected the expression of repetitive DNA elements in innervated and denervated muscles of seven weeks old mice (Fig. 6). The expression of repetitive elements was indistinguishable in CTL and H2A.Z dKO innervated muscle (Fig. 6B). Interestingly, denervation altered the expression of several types of repetitive elements in control muscles. 72 families were up-regulated and 3 were down-regulated 48 hours after denervation (Fig. 6C). These changes were similar in H2A.Z dKO muscles (Fig. 6D, E, Table 3). The MLTH1 and RLTR6 families of repetitive elements were among the most up-regulated by denervation. H2A.Z inactivation affected neither their expression in innervated muscle nor their activation in denervated muscle (Fig. 6E). Taken as a whole, these results indicate that H2A.Z is neither required for normal expression nor for the activation of repetitive DNA elements.

**Figure 6.**
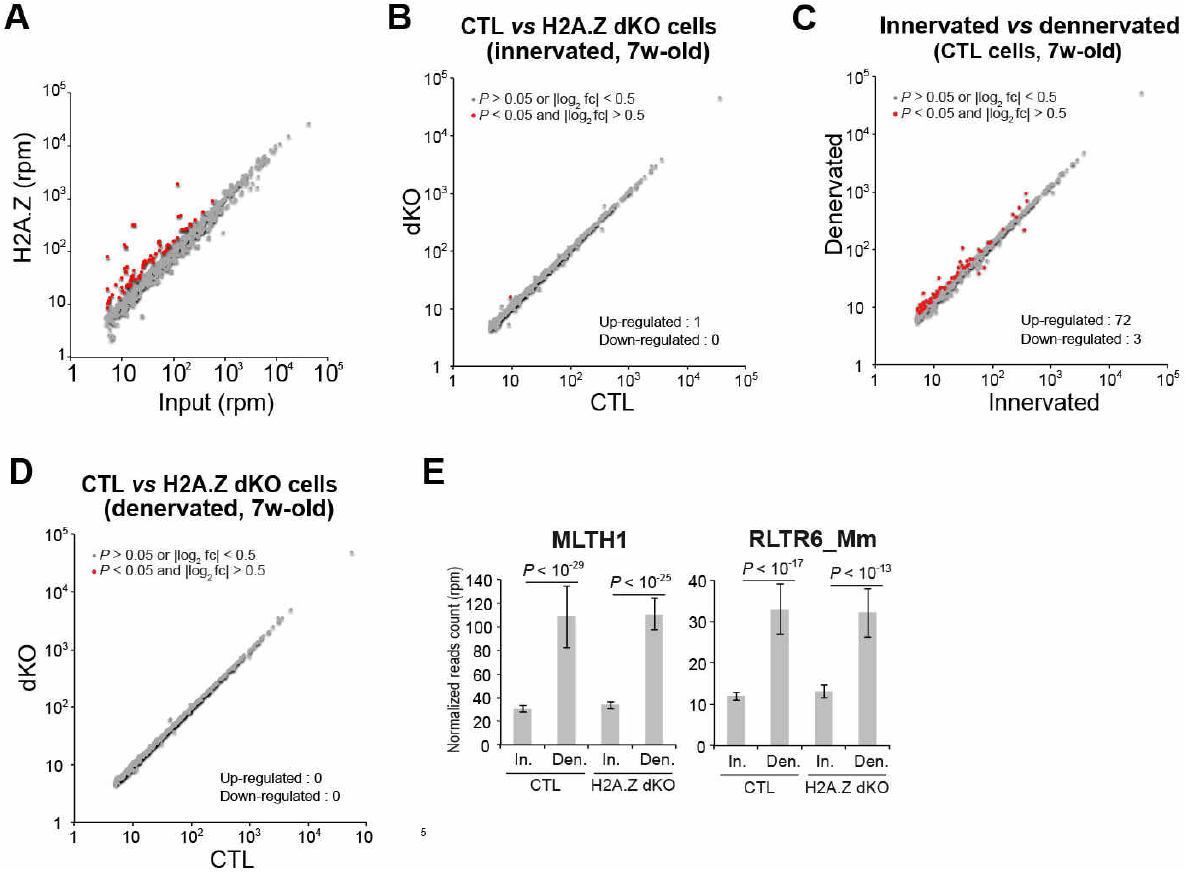
H2A.Z at DNA repetitive elements. **(A)** Scatter plot showing the average enrichment of H2A.Z in repeat families. **(B)** Scatter plots comparing global transcription of repetitive elements between CTL and H2A.Z dKO myofibers in innervated muscle of 7 weeks-old mice. **(C)** Scatter plots comparing global transcription of repetitive elements in innervated and denervated muscle cells (7 weeks-old mice) in the presence of H2A.Z. **(D)** Scatter plots comparing global transcription of repetitive elements between CTL and H2A.Z dKO myofibers in denervated muscle of 7 weeks-old mice. **(E)** Bar graphs representing the expression level of the MLTH1 and RLTR6_Mm retroelements, the two most overexpressed retroelements in response to denervation.

**Table 2.**
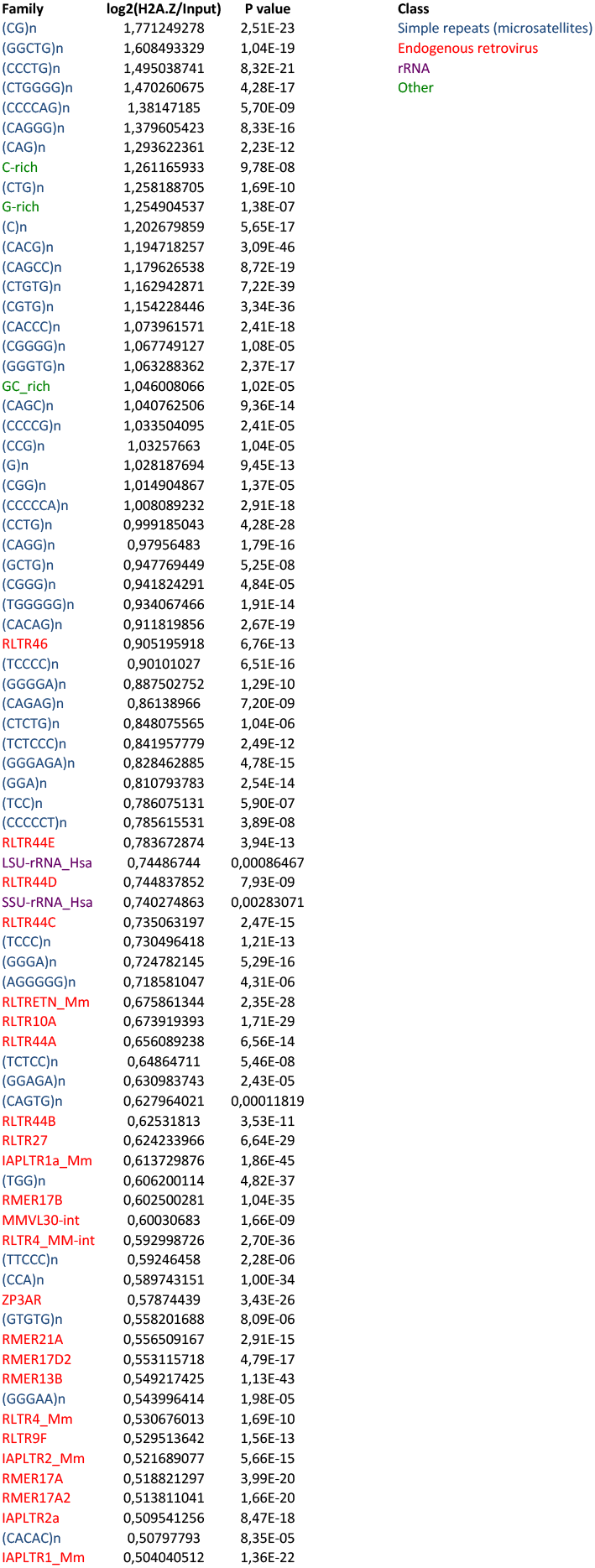
Repetitive elements family enriched of H2A.Z. The majority of significantly H2A.Z-enriched repeats (log2 enrichment > 0.5 and P value < 10-2) belongs to the simple repeats and endogenous retrovirus classes of repetitive elements.

**Table 3.**
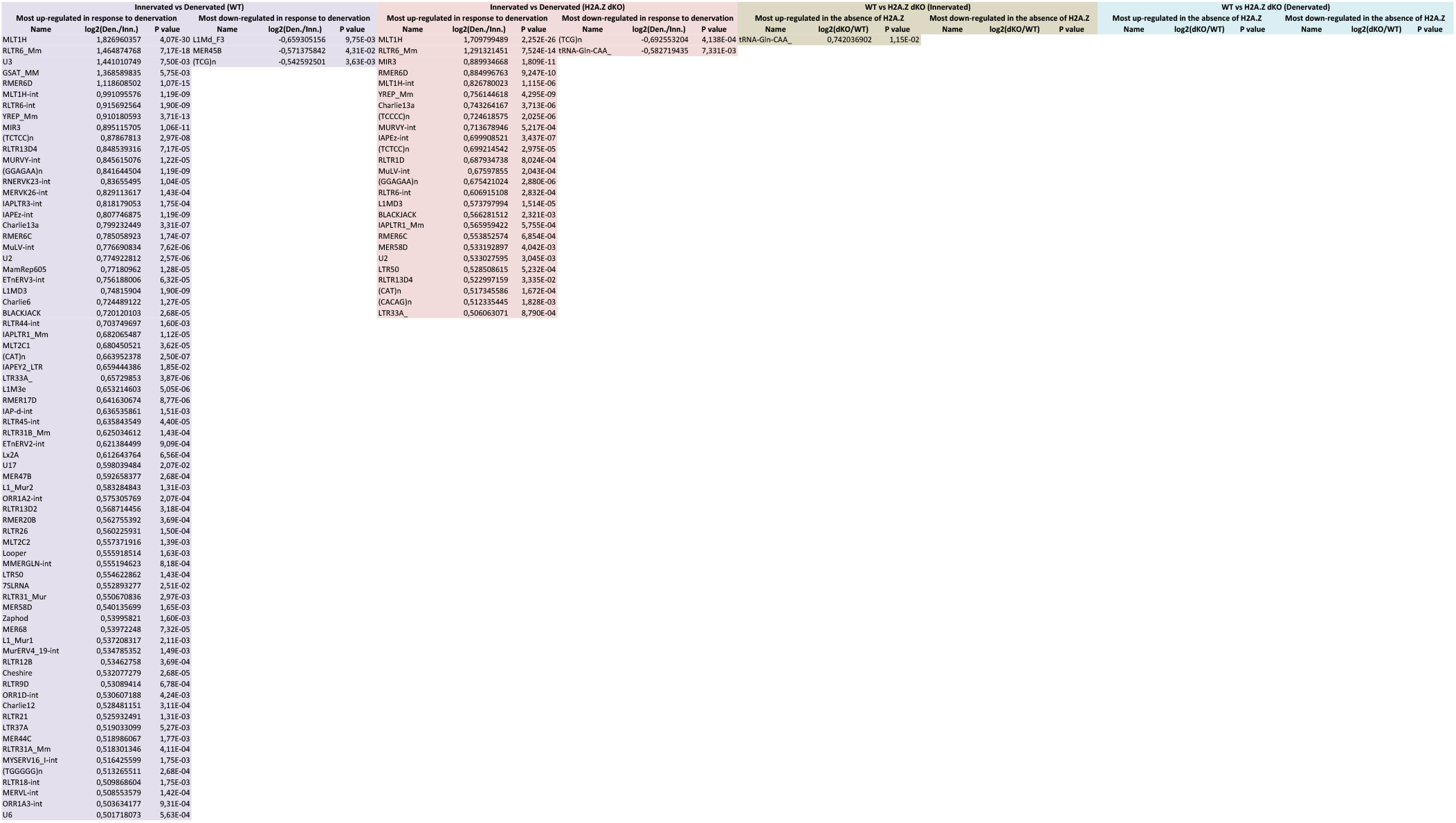
Main down- and up-regulated repetitive DNA element families in innervated and denervated CTL and dKO H2A.Z skeletal muscle from 7 week-old mice.

## DISCUSSION

The goal of this study was to evaluate *in vivo* in post-mitotic tissues the function of H2A.Z in transcription. As model system we have first isolated MEF cells derived from H2AZ.1^flox/flox^:H2AZ.2^flox/flox^ mice. In cycling MEFs, H2A.Z was depleted in less than two days after adeno-Cre infection and many genes were up and down regulated (Fig. 1F and data not shown) according to the literature^25,36,37^. We next sought to avoid a possible influence of the cell cycle on gene expression in the absence of H2A.Z. An *in vivo* approach in skeletal muscle was chosen because it provides access to long lived postmitotic cells in which Cre recombinase expression can be efficiently targeted and where consequences can be analysed several weeks after gene inactivation. In addition, denervation provides a convenient means to rapidly change the transcriptional program of the targeted cells.

ChIP-seq analysis confirmed that in skeletal muscle fibers, H2A.Z was enriched at transcriptional start sites and on regulatory elements as previously described in other systems^15,17,18,23^. To specifically inactivate H2A.Z in the skeletal muscle fibers of H2A.Z-1^flox/flox^:H2A.Z-2^flox/flox^ mice, HSA-Cre mice were used. This Cre driver mouse line is probably the best characterized muscle specific Cre-driver mice and in skeletal muscle they allow to selectively target postmitotic cells^27^.

Skeletal muscles contain a large variety of cell types (fibroblasts, satellite cells macrophages, endothelial cells, …)^34^. The Human Skeletal Actin promoter drives the expression of the Cre recombinase only in differentiated muscle cells. When performing muscle extracts from the whole tissue non muscle cells such as fibroblasts are inevitably present. These cells represent less than 20% of the muscle tissue but they are sufficiently represented to generate detectable expression of inactivated genes in muscle extracts. This is true for H2A.Z dKO muscles and for all muscle specific inactivation of ubiquitously expressed genes^38–42^. By allowing to visualize individual nuclei, immunofluorescence was therefore the best way to evaluate the efficiency of H2A.Z knock-down in muscle fibers. In H2A.Z dKO muscles, H2A.Z staining had already decreased below the detection level by 2 weeks of age and H2A.Z remained undetectable in muscle fibers since then. When analysed seven weeks after birth, H2A.Z dKO mice and H2A.Z dKO muscles did not show any detectable alteration (Fig. 4A, B). Strikingly, RNA-seq analysis revealed no significant effect of H2A.Z depletion on the transcriptomic profile of H2A.Z dKO muscles. Induction of a new transcriptional program by denervation was not perturbed either by the absence of H2A.Z. Moreover, the chromatin accessibility analysed by ATAC-seq did not reveal any differences between control and H2A.Z dKO muscles.

H2A.Z is also preferentially enriched at regulatory elements such as enhancers and CTCF-binding sites, which mark insulator sites in the genome^17^. ATAC-seq results would have shown changes in chromatin accessibility if H2A.Z depletion had perturbed their function. We can therefore conclude that as for proximal promoters regulation, H2A.Z is not required for enhancers and insulators function.

The transcriptional function of H2A.Z is probably not as important as initially thought but it is still an essential protein in cycling cells. It is known that it is enriched at centric and pericentromeric heterochromatin^21,43,44^ and it is also, somehow, involved in mitosis^21^. If H2A.Z has key roles in organizing the centrosome or other chromatin loci crucial for mitosis, they are important enough to lead to cell death when perturbed. The fact that none of these functions are required in post-mitotic cells would explain why H2A.Z is not an essential protein in post-mitotic muscle cells. It would be interesting to investigate the nucleosome composition at TSS in the absence of H2A.Z. Three non-exclusive possibilities would fit with our data. The first, and most likely hypothesis, is that H2A.Z would be replaced by canonical H2A. H2A expression is stronger during S phase^45^, however, low level of H2A expression is probably sufficient in post-mitotic cells. This is the case for histone H4 for which there is no known variant in the mouse genome (to our knowledge H4 has a variant only in hominids^4^). If H4 is expressed at low levels from one of the histone clusters, H2A might be as well. The second possibility is that H2A.Z would be replaced by another H2A variant. There are many H2A variants which can be incorporated in nucleosomes. However, the expression of none of them was upregulated in H2A.Z dKO muscles. The third possibility is that H2A.Z would not be replaced by another histone, thus creating an incomplete nucleosome e.g. a tetrasome at the TSS (see^46^) or even gaps in chromatin if the tetrasome was too unstable. However, ATAC-seq results did not reveal any increase in chromatin accessibility, arguing against this hypothesis. The most likely explanation is therefore that canonical H2A replaces missing H2A.Z molecules, but ruling out the other possibilities will require further in depth investigations.

It has long been thought that since H2A.Z was enriched at TSS, it was playing an active role in transcriptional regulation. Our results showing normal gene expression and regulation in post-mitotic muscle cells challenge this view. Almost all data available so far were obtained using cycling cells and could not rule out the possibility that H2A.Z functions in other processes than transcription. A role in replication or DNA repair could easily explain why H2A.Z total KO in mouse was lethal during early embryonic development. Our results suggest that the requirement of H2A.Z for cellular viability *in vivo* is not linked to transcription. Alternatively, a differential involvement of H2A.Z in transcription in cycling and postmitotic cells could be envisaged. Even though, H2A.Z accumulation at active TSS and regulatory elements with no apparent function in post-mitotic cells would still be puzzling.

If not required for a specific transcriptional function, the accumulation of H2A.Z at TSS and regulatory regions is intriguing. A simple explanation of this phenomenon could be provided by the fact that overall, H2A.Z accumulates at sites where nucleosome turnover is high. H2A.Z would be preferentially incorporated in new nucleosomes at these sites because it is more available than its canonical counterpart mainly expressed during the S phase. H2A.Z would therefore be used to locally reconstitute chromatin and to protect the DNA from damage without playing an active role. Altogether, these results suggest that H2A.Z is rather a marker than an actor of transcriptional activity. Recently, several reports on the essential variant H3.3 similarly claimed that it is dispensable for transcription^47,48^. This raises the question of the role of histone variants in transcriptional regulation as a whole.

## MATERIALS AND METHODS

### Generation of H2A.Z-1/H2A.Z-2 HSA-Cre mice

The H2A.Z mouse line was generated at the “Institut Clinique de la Souris” (ICS-MCI; Illkirch, France). The targeting vector containing a Frt Neomycin resistance cassette and (i) *H2afz*: in which exons 2-4 was flanked by loxP sites or (ii) *H2afv*: in which exons 4-5 was flanked by loxP sites was generated. Each constructs were electroporated into mouse embryonic stem (ES) cells. Targeted ES cells were injected into C57BL/6 blastocysts which were implanted in pseudo-pregnant females. Removal of the Neomycin cassette in the targeted allele was accomplished by crossing the chimeric males giving germline transmission with Flp transgenic females to generate mice with the conditional allele (Fig. 1A and B). Each strain was backcrossed for 10 generations on C57BL/6N mice. Finally, H2A.Z-1^flox/flox^:H2A.Z-2^flox/flox^ mice were obtained by crossing the mice with HSA-Cre transgenic strain^27^ to generate the muscle specific conditional double KO. Mice were genotyped by PCR amplification of genomic DNA extracted from newborn biopsies. A 40 μl volume of extraction buffer (0.5 mg/ml proteinase K, 0.2% SDS, 0.2 M NaCl, 100 mM Tris, 5 mM EDTA, pH 8.0) was added to biopsies and incubated at 54 °C overnight. After DNA precipitation and resuspension in 200 μl of water, 1 μl of DNA was taken as template in a 25 μl PCR reaction using the Go Taq DNA polymerase (Promega) according to the provider’s recommendation. Genotyping was performed with primers listed in supplementary Table 1.

### Cell culture

Mouse embryonic fibroblasts (MEFs) were derived from E13.5 WT and dKO embryos. Heads and internal organs were removed, the torso was minced into chunks of tissue. Cells were immortalized with a retrovirus expressing the large SV40 T antigen and cultured in high glucose Dulbecco’s modified Eagle medium (DMEM), with sodium pyruvate, Glutamax (Gibco), 10% fetal bovine serum (FBS), and penicillin–streptomycin in a humidified incubator at 37°C and a 5% CO2 atmosphere. Immortalized H2A-Z.1^flox/flox^:H2A-Z.2^flox/flox^ MEFs were plated at 30% of confluency and were infected with recombinant adenoviruses encoding either GFP or a Cre-GFP fusion (Ad-CMV-GFP or Ad-CMV-Cre-GFP, Vector Biolabs, Philadelphia, Pennsylvania, USA) to respectively generate control (CTL) and H2A.Z double KO (H2A.Z dKO) cell lines. Viruses were diluted in the culture media for overnight infection. Cells were then washed and were diluted 4 times every 2 days to have them always below a confluence of 70%. Cells were collected every 2 days until 8 days after infection.

### Mice care

Animals were provided with mouse chow and water ad libitum in a restricted-access, specific pathogen–free animal care facility at the Ecole Normale Supérieure of Lyon (Plateau de Biologie Expérimentale de la Souris). All procedures were performed in accordance with national and European legislation on animal experimentation.

### Denervation of hindlimb muscles

Section of the left sciatic nerve was used to induce denervation. Briefly, after the mice were anesthetized with an intraperitoneal injection of ketamine (100 mg.kg^-1^) and xylazine (10 mg.kg^-1^), the sciatic nerve was exposed in the thigh and doubly cut (5 mm apart) just distal to the sciatic notch. Contralateral muscles that were not operated and were used as controls. 48 hours post-denervation, non-denervated and denervated *tibialis anterior* (TA) muscles were collected and snap frozen in liquid nitrogen. All operative procedures were performed using aseptic techniques and according to the ethical committee recommendations (Ceccap-ENS-2014-019).

### Quantitative RT-PCR

For RNA extraction from MEF cells, 1ml of TRI reagent (Sigma) was added to 10 cm cell culture plate that have been washed once before with PBS. Cells are then scrapped and place in a 1.5 ml tube and processed following provider’s instructions. To extract total RNA from muscles, 500 μl of TRI Reagent (Sigma) was added to individual frozen Tibialis Anterior in tubes containing ceramics beads (Lysing Matrix D – MP biomedicals) for homogenization in a PreCellys (Bertin Technologies) (6500 RPM, 3 × 10 s). After centrifugation, TRI Reagent was removed and beads were washed once with 500 μl of TRI reagent. Total RNA was extracted following provider’s instruction. To generate cDNA, total RNA was treated with DNase (Ambion) and reverse transcribed with RevertAid H Minus Reverse Transcriptase (ThermoFisher) primed with random hexamers. RT–qPCR was performed using SYBR Green Mastermix (Qiagen) in the CFX-connect system (Bio-Rad). Relative expression levels were normalized to *GusB* and *Rpl41* housekeeping genes expression using the ΔΔ*C_t_* method. Primers are listed in Supplementary Table 2.

### RNA-seq

Libraries of template molecules suitable for strand-specific high-throughput DNA sequencing were created using a TruSeq Stranded Total RNA with Ribo-Zero Gold Prep Kit (RS-122-2301; Illumina) as previously described^49^. The libraries were sequenced on Illumina Hiseq 4000 sequencer as single-end 50 bp reads following Illumina’s instructions. Image analysis and base calling were performed using RTA 2.7.3 and bcl2fastq 2.17.1.14. Adapter dimer reads were removed using DimerRemover v0.9.2. Reads were mapped to the mouse genome (mm9) using Tophat^48^ v2.0.14 and Bowtie^50^ v2-2.1.0. Quantification of gene expression was performed using HTSeq^51^ v0.6.1 and gene annotations from Ensembl release 67.

### Nuclei preparation

Muscles of individual hind limbs were collected and put in 4 ml of cold buffer A (300 mM sucrose, 10 mM NaCl, 1.5 mM MgCl2, 15 mM Tris HCl pH 8.0 and protease inhibitor cocktail (Sigma)). Muscles were minced with scissors and transfered in a dounce homogenizer where 50 strokes of the loose pestle were slowly applied.

The extract was then passed through a 100 μm cell strainer to remove myofibrils debris and placed back into a clean cold dounce homogenizer. Nonidet P-40 was then added to the extract to a final concentration of 0.5%, which was then incubated for 15 min at 4°C. Subsequently, 50 strokes with the tight pestle were applied. The nuclei suspension was finally clarified through a 40 μm cell strainer, centrifuged at 2000 g for 10 min at 4°C and washed in cold buffer A. Purified nuclei were resuspended in a small volume of buffer A for further ChIP-seq, ATAC-seq or immunoblot analysis.

### ChIP-seq

Purified nuclei were crosslinked with 0.2% formaldehyde for 1 min before quenching with 125 mM glycine for 5 min at room temperature. The buffer was then changed to 300 mM NaCl, 300 mM sucrose 0.1% NP-40, 2 mM CaCl2, 15 mM Tris/HCl pH 8.0 following a centrifugation at 2000 g for 10 min. MNase (Roche diagnostic) was then added to the nuclei and incubated at 37°C for a period of time long enough to generate a majority of mono- and di-nucleosomes. The reaction was stopped by increasing EDTA concentration to 10 mM. After a 5 min centrifugation at 20 000 g, 20 μg of chromatin from the supernatant were incubated overnight at 4°C with an immunopurified highly specific polyclonal anti-H2A.Z rabbit antibody generated in-house^15^, followed by 2 hours incubation with Protein A dynabeads (Invitrogen). Immunoprecipitated material was sequentially washed with buffer containing low salt (150 mM NaCl), high salt (500 mM NaCl) and finally a high stringency wash (500 mM NaCl, 0.25 M LiCl and 1% sodium deoxycholate) in 10 mM Tris/HCl pH 8, 1 mM EDTA buffer. DNA was released from immunoprecipitated complexes by overnight incubation at 65°C and finally isolated over Qiagen PCR purification columns.

Libraries were prepared using the Diagenode MicroPlex Library Preparation kit v2, and sequenced on Illumina Hiseq 4000 sequencer as single-end 50 bp reads following Illumina’s instructions. Image analysis and base calling were performed using RTA 2.7.3 and bcl2fastq 2.17.1.14. Adapter dimer reads were removed using DimerRemover v0.9.2. Reads were mapped to the mouse genome (mm9) using Bowtie^2^ v1.0.0 with the following arguments: -m 1 --strata --best -y -S –l 40 -p 2.

### ATAC-seq

Purified nuclei are quantified using DAPI and 50 000 of them are taken to performed the ATAC-seq protocol as described in Buenrostro et al^52^. Nextera adapters were trimmed from reads with Trimmomatic^53^. Remaining paired reads longer than 20bp were mapped to the mouse genome (mm9) using Bowtie2^2^, using the local alignment mode (--local) and a maximum fragment length set to 2000bp (--maxins).

### Immunoblot

Chromatin preparations were separated on a 18% SDS-PAGE and transferred onto polyvinylidene fluoride (PVDF) Immobilon-P membranes (Millipore). Immunoblots were performed with enhanced chemiluminescence (ECL) PLUS reagent (GE Healthcare) according to the manufacturer’s instructions. Antibodies used were as follows: rabbit monoclonal anti-Histone H4 pan (04-858, Merck Millipore); rabbit polyclonal anti-H3 (#61277); mouse monoclonal anti-α-Tubulin (T6074, Sigma); the rabbit polyclonal anti-H2A.Z^15^.

### Histology

Hind limb muscles surrounding the tibial bone (Gastrocnemius, Plantaris, Soleus (GPS), TA and Extensor Digitorum Longus (EDL) were collected from 7 weeks-old CTL and H2A.Z dKO mice, frozen in isopentane cooled on dry ice, and cross-sectioned at 10 μm thickness in a cryostat. Transverse sections were stained with Hematoxylin and Eosin for immunohistochemistry and with Wheat Germ Agglutinin (WGA) / DAPI for immunofluorescence, analysed using an Axio Scan.Z1 slide scanner (Zeiss Microscopy). Morphometrics analysis were performed with the ImageJ software.

### Immunofluorescence

Cross sections were rehydrated in PBS, fixed with 4% PFA for 10 min, permeabilized in 1X PBS, 0.1% Triton for 30 min at room temperature then saturated in 1X PBS 1% BSA 1% normal goat serum for 1 hour. Staining with green WGA (W11261, Molecular Probes) and DAPI were performed overnight at 4°C. Coverslips were then mounted with Vectashield (Vector Laboratories) and sealed with nail polish.

For myofibers staining, an EDL muscle was collected and fixed in 4% paraformaldehyde 10 min for tissue dissection and staining. Muscles were slowly teased with fine forceps to isolate individual fibers and small fiber bundles. Fibers were permeabilized in 1X PBS, 0.1% Triton for 30 min at room temperature, saturated in 1X PBS 1% BSA for 1 hour before overnight incubation with the primary antibody (home-made anti-H2A.Z 1/100) diluted in blocking buffer. After washing, fibers were incubated with a secondary antibody conjugated with FITC (Molecular probes). DNA and neuromuscular junctions were respectively stained with DAPI (2 μg/ml) and α-bungarotoxin Alexa-555 (Molecular probes) for 2 hours at room temperature. After washing, fibers were mounted on glass slide with Vectashield (Vector Laboratories) and analysed using a confocal laser scanning microscope (Leica SP5). Images were processed using ImageJ software.

### Computational analyses

Repeat analyses of RNA-seq and ChIP-seq datasets were performed as previously described^1^. Processed datasets were restricted to repeat families with more than 500 mapped reads per ChIP sample or more than 5 reads per million mapped reads per RNA sample to avoid over- or underestimating fold enrichments due to low sequence representation.

Heatmaps and quantitative analysis of the ChIP-seq data were performed using seqMINER (http://bips.u-strasbg.fr/seqminer/). As reference coordinates, we used the Ensembl 67 database of the mouse genome (mm9).

For ATAC-seq data not properly paired, supplementary alignments and putative PCR duplicates were removed^54–56^. Enriched regions were identified with MACS2^57^ at q value 0,01, with either a wide or narrow assumption of how accessibility is inferred from fragments ends: a wide model was a half nucleosome length (73bp) centered on the fragment ends (ie --nomodel --shift −37 -- extsize 73), these ends having been shifted 4bp forward for (+) reads and 5bp backward for (-) reads, in order to represent the center of the tn5 transposon binding event, as suggested^58^; the narrow model was the tn5 transposon occupancy only (ie --nomodel --shift −5 --extsize 9). Mitochondrial peaks and peaks overlapping blacklisted regions were filtered out^59^. In order to compare accessibility between both conditions, we focused on the union of their peaks, analysing those called with the wide or narrow model separately. As an example, for the peaks produced with the wide calling model, we first defined true enriched regions in each condition as MACS2 peaks shared by the three replicates, taking the union of the intervals, if overlapped by at least 25% of their length; we then merged these true enriched regions from both conditions, taking the union of regions overlapping by more than 50% of their length, or keeping not overlapped regions with their boundaries unchanged^60^. We quantified the fragments ends tn5-shifted and 37bp extended falling within these 52142 union intervals. Differential peak enrichment was then computed using DESeq2^61^, Heatmaps were performed using deeptools2^62^, plotting the means of the fold enrichment files generated by MACS2 from the three replicates for each condition, using the narrow peak calling, the TSSs being sorted either by the RNA-seq expression data from the CTL condition, or by the CTL ATAC-seq TSS enrichment.

### Data access

The ChIP-seq and RNA-seq datasets have been deposited in the Gene Expression Omnibus (GEO; http://www.ncbi.nlm.nih.gov/geo/) under the accession number GSE111576.

## SUPPLEMENTARY DATA

Supplementary Data are available online.

## ACKNOWLEDGEMENTS

Animal breeding and H2A.Z muscle-specific inactivation were performed at the animal facility (PBES) of the research federation SFR Biosciences (UMS3444). Microscopy was performed on the microscopy facilities of SFR Biosciences (PLATIM, UMS3444) and SFR Santé Lyon-Est (CIQLE, UMS 3453).

## AUTHOR CONTRIBUTIONS

L.S., S.D., A.H., E.B., N.L. conceived the research. L.R. and D.D. participated to the generation of KO mice, E.B. performed the conditional KO, MEF cells and mouse breeding. E.B., N.L. performed cell biology, RT-QPCR, immunofluorescences, histology, nuclei isolation and biochemistry analysis. K.P. performed ChIP experiments. I.S., C.P., T.S. performed bioinformatics analysis. E.B., N.L., L.S., S.D., A.H. wrote the paper with input from all authors.

