## Supplementary for "H2A.Z is dispensable for both basal and activated transcription in post-mitotic mouse muscles"

**SUPPLEMENTARY INFORMATIONS**

**Supplementary materials and methods**

**Histology analysis**

Serial cross-sectioned (10µm thick) were collected along the entire length of embedded muscle tissues. Four consecutive slices were stained for following histological and histochemical analysis: HE, Gomori's Trichrome (TRICH), cytochrome oxidase (COX), succinate dehydrogenase (SDH), and reduced nicotinamide adenine dinucleotide tetrazolium reductase (NADH-TR). Staining was done according to standard protocols.

**Supplementary Figures**

**Supplementary Figure 1:** Read sum up obtain from the RNA-seq of H2A.Z dKO muscle obtain around each H2A.Z genes. Note that reads are present only in the beginning of each H2A.Z genes and not in the regions located between loxP sites (end of each genes).

**Supplementary Figure 2:** RT-qPCR analyses of MyoD (**A**) and Myogenin (**B**) after denervation of TA muscle in presence (CTL) or absence of H2A.Z (H2A.Z dKO).

**Supplementary Figure 3:** Phenotypic characterisation of control and H2A.Z dKO mice in innervated and denervated conditions after 48h. **(A)** Transverse sections of TA muscle were histochemically stained with Haematoxylin Eosin (HE), Gomori's Trichrome (TRICH), SDH-CoxH and NADH activity. (scale bar 500µm) **(B)** Zoom in of supplementary figure 1A with the same staining. (scale bar 100µm)

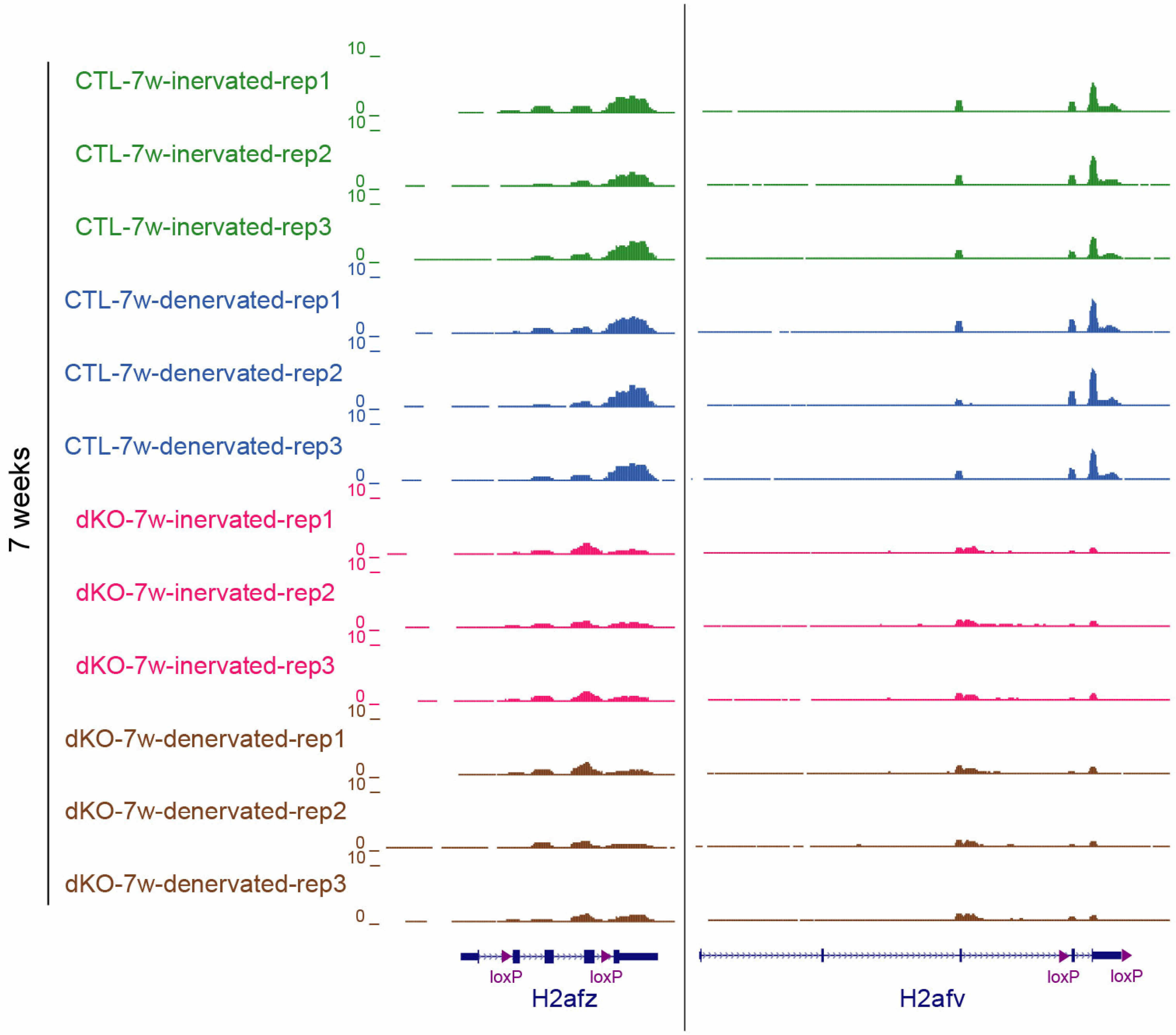


Supplementary Figure 1

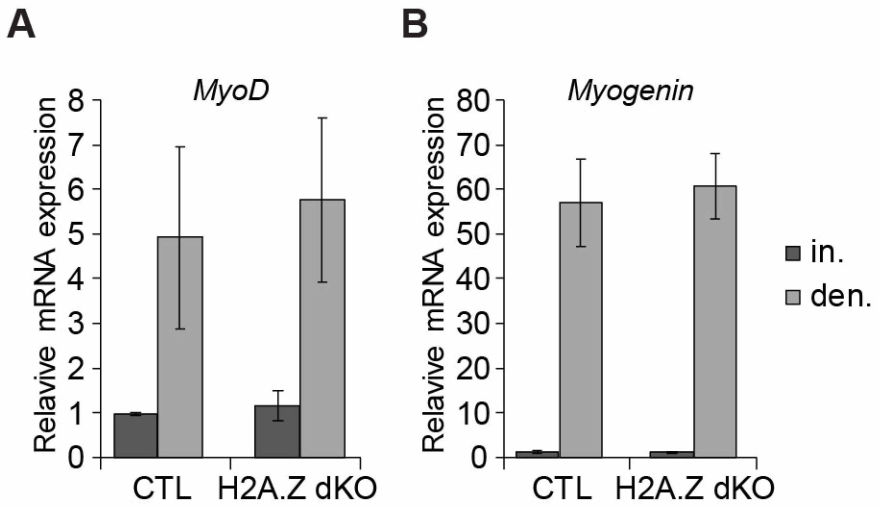


Supplementary Figure 2

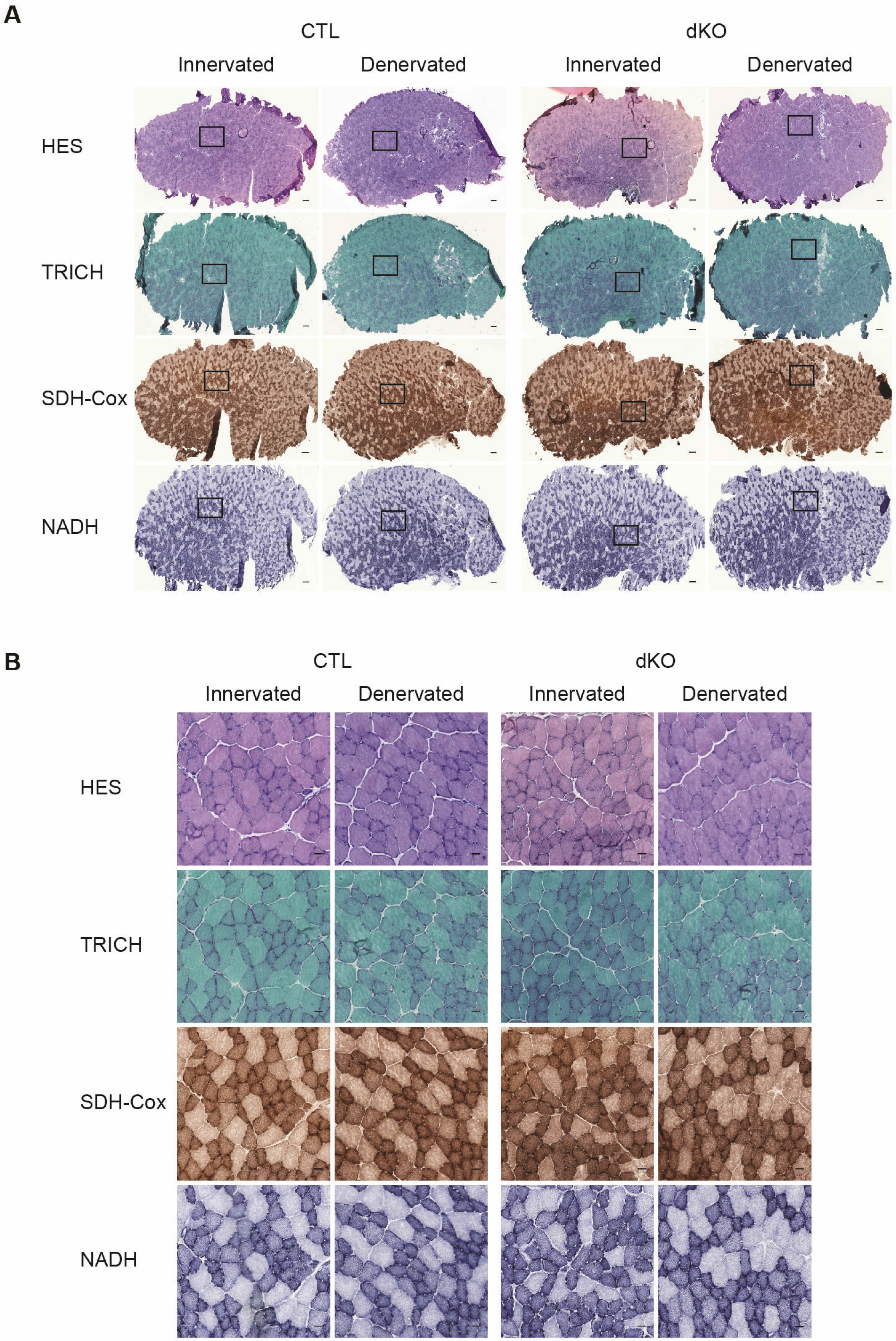


Supplementary Figure 3
